# Comparative genetic, biochemical, and biophysical analyses of the four *E. coli* ABCF paralogs support distinct functions related to mRNA translation

**DOI:** 10.1101/2023.06.11.543863

**Authors:** Farès Ousalem, Shikha Singh, Nevette A. Bailey, Kam-Ho Wong, Lingwei Zhu, Matthew J. Neky, Cosmas Sibindi, Jingyi Fei, Ruben L. Gonzalez, Grégory Boël, John F. Hunt

## Abstract

Multiple paralogous ABCF ATPases are encoded in most genomes, but the physiological functions remain unknown for most of them. We herein compare the four *Escherichia coli* K12 ABCFs – EttA, Uup, YbiT, and YheS – using assays previously employed to demonstrate EttA gates the first step of polypeptide elongation on the ribosome dependent on ATP/ADP ratio. A Δ*uup* knockout, like Δ*ettA*, exhibits strongly reduced fitness when growth is restarted from long-term stationary phase, but neither Δ*ybiT* nor Δ*yheS* exhibits this phenotype. All four proteins nonetheless functionally interact with ribosomes based on *in vitro* translation and single-molecule fluorescence resonance energy transfer experiments employing variants harboring glutamate-to-glutamine active-site mutations (EQ_2_) that trap them in the ATP-bound conformation. These variants all strongly stabilize the same global conformational state of a ribosomal elongation complex harboring deacylated tRNA^Val^ in the P site. However, EQ_2_-Uup uniquely exchanges on/off the ribosome on a second timescale, while EQ_2_-YheS-bound ribosomes uniquely sample alternative global conformations. At sub-micromolar concentrations, EQ_2_-EttA and EQ_2_-YbiT fully inhibit *in vitro* translation of an mRNA encoding luciferase, while EQ_2_-Uup and EQ_2_-YheS only partially inhibit it at ~10-fold higher concentrations. Moreover, tripeptide synthesis reactions are not inhibited by EQ_2_-Uup or EQ_2_-YheS, while EQ_2_-YbiT inhibits synthesis of both peptide bonds and EQ_2_-EttA specifically traps ribosomes after synthesis of the first peptide bond. These results support the four *E. coli* ABCF paralogs all having different activities on translating ribosomes, and they suggest that there remains a substantial amount of functionally uncharacterized “dark matter” involved in mRNA translation.

---

<u>A</u>TP-<u>b</u>inding <u>c</u>assette (ABC) superfamily proteins employ a pair of stereotyped ATPase domains to perform mechanical and regulatory activities ^1–6^. Although most domains in this superfamily are components of ATP-powered transmembrane transporters, a sizable fraction of ABC domains are found in soluble protein complexes that mediate activities unrelated to transport, ranging from DNA repair to regulation of mRNA translation into protein ^7,8^. The ABCF family is the most widely distributed and diverse class of soluble proteins within the ABC superfamily ^7,9–11^. The function has only been characterized in detail for a relatively small number of ABCF paralog groups, including that containing the *Escherichia. coli* protein EttA ^12–14^ and a series of proteins called <u>A</u>ntibiotic <u>R</u>esistance <u>E</u>lements (AREs) that confer ^8,15–33^ or regulate ^34,35^ resistance to ribosome-targeting antibiotics. The physiological functions of all of the other ABCF paralog groups remain imprecisely defined or entirely unknown, despite their very broad phylogenetic prevalence ^7,10,11,13,36^ and their involvement in a variety of medically important processes. For example, human ABCF paralogs have been identified in a variety of genome-scale screens for factors modulating processes ranging from inflammation ^37–39^ and innate immunity ^40^ to responsiveness to cancer chemotherapy ^41–49^, but the biochemical mechanisms underlying these activities is unclear.

ABCF proteins have two tandem ABC ATPase domains ~230 residues in length (PFAM ^12,15–20^ domain PF00005) flanking a domain ~80 residues in length that we dubbed the P-site tRNA interaction motif ^13^ (PtIM – PFAM domain PF12848). Multiple ABCF paralogs likely to have different molecular functions are encoded in the genomes of the most widely studied model organisms ^7,13^. *Saccharomyces cerevisiae* encodes two, *Arabidopsis thaliana* encodes five, humans encode three ^9,50^, and *E. coli* encodes four (EttA, Uup, YbiT, and YheS). ABCF proteins belonging to several dozen different paralog groups likely to be functionally distinct are encoded in eubacterial genomes, and the vast majority of them encode at least one ^36^.

We previously demonstrated that *E. coli* EttA binds in the tRNA exit (E) site of a ribosomal 70S initiation complex (IC), where its PtIM contacts the fMet-tRNA^fMet^ in the peptidyl-tRNA binding (P) site ^12,14,51^ and gates synthesis of the first peptide bond by the 70S IC ^13^. A super stoichiometric ratio of ADP over ATP causes EttA to inhibit formation of the first peptide bond, but this inhibition is relieved when the concentration of ATP rises above that of ADP ^13^. Binding of ATP to EttA has been shown to then promote synthesis of the first peptide bond by the 70S IC. Introducing hydrolysis-deficient glutamate-to-glutamine (EQ_2_) mutations in the active sites of EttA trap it in the ATP-bound state, resulting in the formation of a stable EttA-bound ribosomal <u>pre</u>-translocation (PRE) complex ^12^ in which the first peptide bond is formed, but the resulting dipeptide product remains bound to the tRNA in the aminoacyl-tRNA-binding (A) site on the ribosome ^12,13^. The enzymological and mechanochemical properties of the EQ_2_ mutations are described in detail in the accompanying paper ^51^. The EQ_2_-EttA•ATP_2_-bound PRE complex ^12,13,51^ is trapped in a specific conformation called Global State 1 (GS1) or Macrostate I (MS-I), which has the ribosomal subunits in their non-rotated orientation ^52^ and the P and A site tRNAs in their classical configurations. The uL1 stalk on the large (50S) ribosomal subunit is in an open conformation, although its interactions with EttA in the E site shift its conformation relative to the standard conformation in a PRE complex in GS1 in the absence of an ABCF ^14,51,53^. These results led us to hypothesize that EttA prevents commitment of energetic resources to starting synthesis of proteins that cannot be completed in energy-depleted cells, which are characterized by an elevated ADP:ATP ratio ^54,55^. This hypothesis is supported by our observation that knockout of the *ettA* gene severely reduces the competitive fitness of energy-depleted *E. coli* cells when reinitiating growth after long-term residency in stationary phase ^13^.

While precise biochemical and physiological functions have not been defined for any ABCF protein other than EttA and the AREs, experimental observations suggest that proteins belonging to several ABCF paralog groups interact with ribosomes and play some role related to mRNA translation. Defective ribosome biogenesis is observed when the activity of one of the two *Saccharomyces* ABCF paralogs is reduced ^56^, and this protein was also found bound to the 50S subunit in the cryogenic electron microscopy (cryo-EM) structure ^28–31^ of a ribosome-associated <u>q</u>uality <u>c</u>ontrol (RQC) complex ^57^. The other *Saccharomyces* paralog contains an N-terminal extension required for the activity of a kinase that phosphorylates eukaryotic translation initiation factor 2 alpha (eIF2α) in response to amino-acid starvation ^58^. One of the human ABCF paralogs interacts with eIF2α ^59,60^, which recruits the initiator tRNA to the small (40S) subunit of the ribosome, and this paralog attenuates translation initiation and reduces the stringency of start-site selection when locked in the ATP-bound state by EQ_2_ mutations equivalent to those we used to characterize EttA ^61,62^.

Our cryo-EM structure of ribosome-bound *E. coli* EttA harboring such mutations ^12^ was used to explain the influence of sequence polymorphisms in the PtIM of one ARE protein, Vga(A) on differences in the resistance of *Staphylococcus aureus* RN4220 to a panel of structurally diverse antibiotics that target the peptidyl-transferase center (PTC) of the ribosome ^25^. The authors of this study concluded that Vga(A) is likely to displace these antibiotics from the PTC by binding to ribosomes in a similar geometry to EttA. Their mechanistic hypothesis was supported by subsequent *in vitro* studies of ribosome interaction with one of those antibiotics ^33^, while their inference regarding the binding geometry has been confirmed by structural studies of eight different ARE proteins ^27–32^.

Relatively little prior research has been reported on the *E. coli* ABCF paralogs other than EttA. The EttA, Uup, YbiT, and YheS proteins are expressed at levels of ~2, ~0.1, ~0.8, and ~0.2 µM, respectively, under baseline growth conditions in chemically defined media ^63,64^. Knockout of the *ybiT* gene in *Erwinia chrysanthemi* AC4150 (87% identical to *E. coli* YbiT) reduced competitive fitness when infecting plants ^65^. This result was assumed to reflect loss of antibiotic efflux activity, but no related biochemical experiments were performed. The *E. coli uup* gene was first identified in a screen for mutations that enhance the frequency of precise excision of transposons ^66–68^. The Uup protein was later purified ^69^ and demonstrated to have non-specific DNA-binding activity enhanced by the presence of an ~80 residue C-terminal extension belonging to the PF16326 domain family ^70–72^. This extension is required for the effect of Uup on transposon excision *in vivo*, but the purified protein has not been demonstrated to have any enzymatic activity related to transposon excision. Uup has also been demonstrated to partially complement a ribosome biogenesis defect caused by knocking out the *bipA* gene ^36,73^, while all four *E. coli* paralogs have been shown to inhibit protein synthesis *in vivo* when expressed harboring hydrolysis-deficient EQ_2_ mutations equivalent to those we used with EttA ^36^.

Genome-scale analyses provide some information on the biological associations of the *E. coli* ABCF paralogs. The transcription of all four is correlated with that of ribosomal proteins, translation factors, and enzymes that modify the translation apparatus ^74–76^. High-throughput pull-down experiments in *E. coli* ^76,77^ have identified complexes containing Uup and either translation elongation factor (EF) Tu or translation initiation factor (IF) 2, which is not homologous to eIF2, but performs a similar function promoting the binding of fMet-tRNA^fMet^ to the small (30S) subunit of the ribosome. In many eubacteria, the *uup* gene is located immediately downstream of the *rlmL* gene encoding a ribosomal methyltransferase, and these two genes form a co-transcribed operon in *E. coli* ^78,79^. The *yheS* gene in *E. coli* forms a co-transcribed operon with the *yheT* and yheU genes^76,77^, which are frequently located adjacent to the *yheS* gene in other organisms and respectively encode a putative esterase of unknown substrate specificity and a protein of unknown function^78,79^. The *ettA* and *ybiT* genes are frequently located immediately adjacent to divergently transcribed genes for enzymes related to peptidoglycan metabolism (*slt* and *ybiS*, respectively). Knockout of *ettA* modestly increases the permeability of the cell envelope for *E. coli* growing on low-salt medium, but the mechanism underlying this effect is unknown ^80^.

We set out to clarify the range of molecular functions performed by ABCF proteins by systematically comparing the properties of the four *E. coli* K12 paralogs (**Fig. 1a**) using the key genetic, biochemical, and biophysical assays that we used to elucidate the function of EttA ^13^. These proteins all share ~30% pairwise sequence identity (**Fig. 1b** in ref. ^51^, although EttA has slightly stronger homology to Uup than to YbiT or YheS). EttA and Uup orthologs are encoded in ~50% of eubacterial genomes, while YbiT and YheS orthologs are encoded in ~40% and ~30% of eubacterial genomes, respectively. Our results indicate that the *E. coli* ABCF paralogs have diverse biochemical and physiological functions and that at least some of these functions involve novel interactions with translating ribosomes. These results, in conjunction with the cryo-EM structures of 70S ribosome complexes of all four paralogs reported in the accompanying paper ^51^, significantly extend understanding of the biological properties of ABCF paralogs.

**Figure 1.**
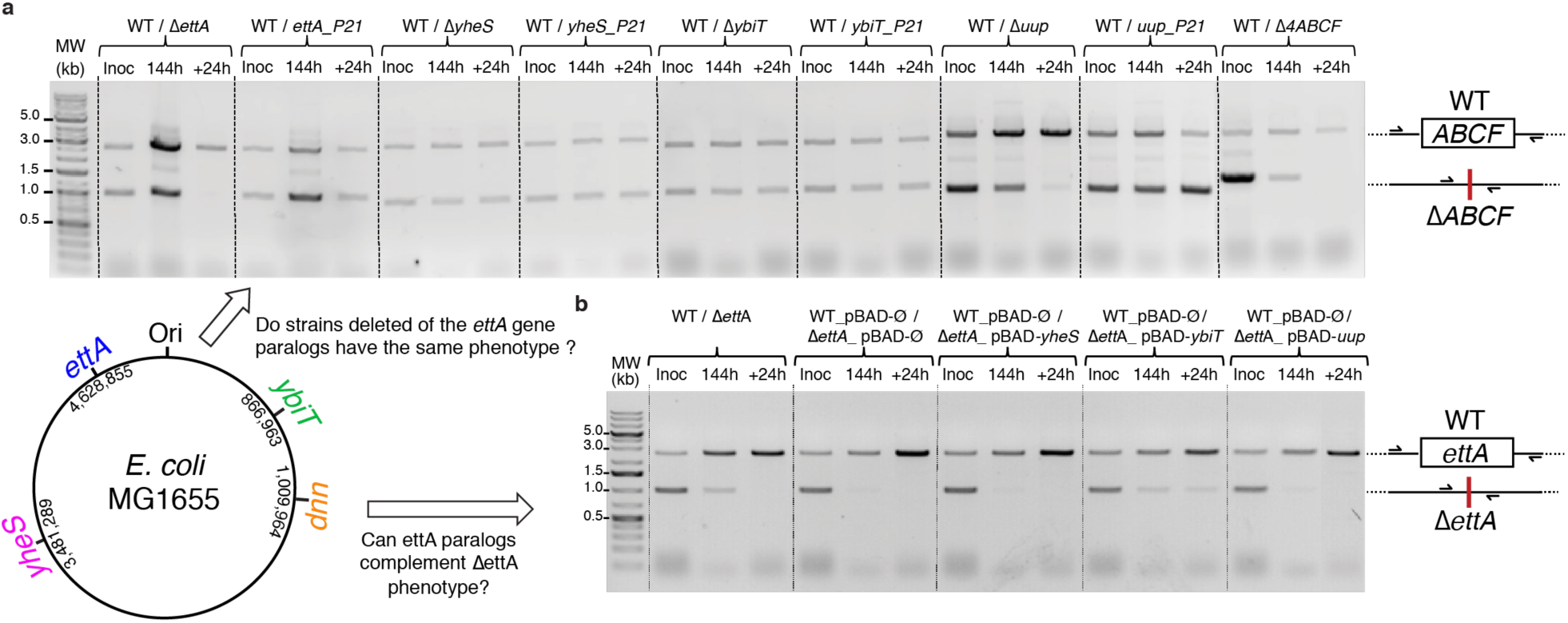
Comparison of the effects of deleting each of the four E. coli ABCF paralogs on fitness in long-term stationary phase. A map of the *E. coli* MG1566 chromosome with the locations of the four ABCF proteins (*ybit*, *uup*, *yheS* and *ettA*) is shown at lower left. PCR assays were performed to amplify the DNA from mixed cultures growing at 37 °C in unbuffered LB in order to quantify the relative levels of WT cells *versus* cells harboring a deletion of one of the ABCF genes. The primers flanked the site of the deleted gene so that the size of the DNA fragment amplified from WT cells is larger than that from cells harboring the deletion. **(a)** Mixed cultures containing isogenic WT and ABCF deletion strains were assayed immediately after inoculating independently grown overnight cultures of the two strains into fresh medium, after six days of growth in that medium, and one day after reinoculating that mixed culture into fresh medium. Experiments were performed on the indicated deletion strains as well as the corresponding complemented deletion strains with the deleted ABCF gene inserted at the P21 locus (“_P21” constructs). Approximately equal numbers of cells from the two strains were mixed together to initiate the assay, based on measurement of the OD_600nm_ of the independently grown overnight cultures. **(b)** Equivalent PCR fitness assays were performed on mixed cultures of the WT strain and a Δ*ettA* strains transformed with either an empty plasmid (pBAD-Ø) or a plasmid expressing one of the three *ettA* paralogues (pBAD-*yheS*, pBAD-*ybiT*, or pBAD-*uup*).

## RESULTS

### Uup, YbiT, and YheS have distinct physiological functions from EttA

We constructed *E. coli* MG1655 strains with the four ABCF paralogs knocked out ^81,82^ individually or simultaneously, and we evaluated the competitive fitness of two independent isolates of each of these strains compared to the parental WT strain when restarting growth in fresh medium after six days of growth in LB at 37 °C. As we demonstrated previously ^13^, the relative proportion of the Δ*ettA* strain declines in mixed cultures with WT MG1655 during the six-day growth period, and the mutant cells are completely lost following one day of growth after reinoculation into fresh medium (**Fig. 1b-c**). Reinsertion of the WT *ettA* gene under endogenous promoter control at an alternative site in the *E. coli* chromosome (*i.e.*, the P21 locus) fully complements the fitness defect in the Δ*ettA* strain (left in **Fig. 1c**), as previously shown for the WT *ettA* gene expressed under arabinose promoter control from a plasmid ^13^. Similar to previously reported results ^83^, the Δ*uup* strain exhibits an equivalent loss of competitive fitness as the Δ*ettA* strain in this assay, as does the ABCF tetra-knockout strain (Δ*ettA*, Δ*uup*, Δ*ybiT*, Δ*yheS*), and reinsertion of the WT *uup* gene under endogenous promoter control at the P21 locus fully complements the fitness defect observed in the Δ*uup* strain. In contrast, the Δ*ybiT* and Δ*yheS* strains show no loss in competitive fitness in this assay compared to the WT strain (**Fig. 1c**), indicating these genes/proteins have distinct physiological functions.

We also tested whether the fitness defect in the Δ*ettA* strain can be complemented by overexpressing each of the three other *E. coli* ABCF paralogs under arabinose promoter control from a multicopy plasmid (**Fig. 1b**). Neither Uup nor YheS overexpression mitigated the Δ*ettA* fitness defect, but YbiT overexpression showed partial complementation when restarting growth in fresh medium after six days of growth in a mixed culture. The observation that Uup overexpression cannot complement the Δ*ettA* defect even though it shows a similar phenotype, while YbiT overexpression can partially complement it even though it does not show a related phenotype, demonstrates that all four of the *E. coli* ABCF paralogs have different physiological activities that presumably reflect differences in their biochemical functions.

### EQ_2_ mutants of the four E. coli ABCF paralogs inhibit cell growth in logarithmic growth phase

For each of the four *E. coli* ABCF proteins, we constructed hydrolysis-deficient EQ_2_ mutants in which both ABC domains have isosteric glutamine residues substituted for the catalytic glutamate residues that activate water for hydrolysis of ATP. We have used equivalent mutations in the isolated ABC domains of two bacterial transmembrane transporters to demonstrate that the binding of two Mg-ATP molecules to a pair of ABC domains drives formation of an “ATP-sandwich” dimer in which the two domains make extensive packing contacts surrounding the Mg-ATP molecules bound at their mutual interface ^84,85^, and a wide variety of subsequent studies support the ATP-driven formation of this complex being the mechanochemical powerstroke in most ABC superfamily proteins ^86–96^. Equivalent mutations trap a wide variety of ABC superfamily proteins ^97–101^, including EttA ^13^ and seven different ARE proteins^27–32^, in equivalent ATP-bound conformations, and the accompanying paper reporting cryo-EM structures of 70S ribosomal complexes of the four *E. coli* ABCF proteins ^51^ demonstrates that EQ_2_ mutations trap all of them in closely related ATP-bound conformations. We previously demonstrated ^13^ that EQ_2_-EttA strongly inhibits cell growth when modestly overexpressed *in vivo*, whereas expression of WT EttA at an equivalent level has no effect on cell growth (**Fig. 2a**). Biochemical and biophysical studies demonstrated that the influence on cell growth is attributable to inhibition of protein synthesis by ATP-bound EQ_2_-EttA ^12,13^.

**Figure 2.**
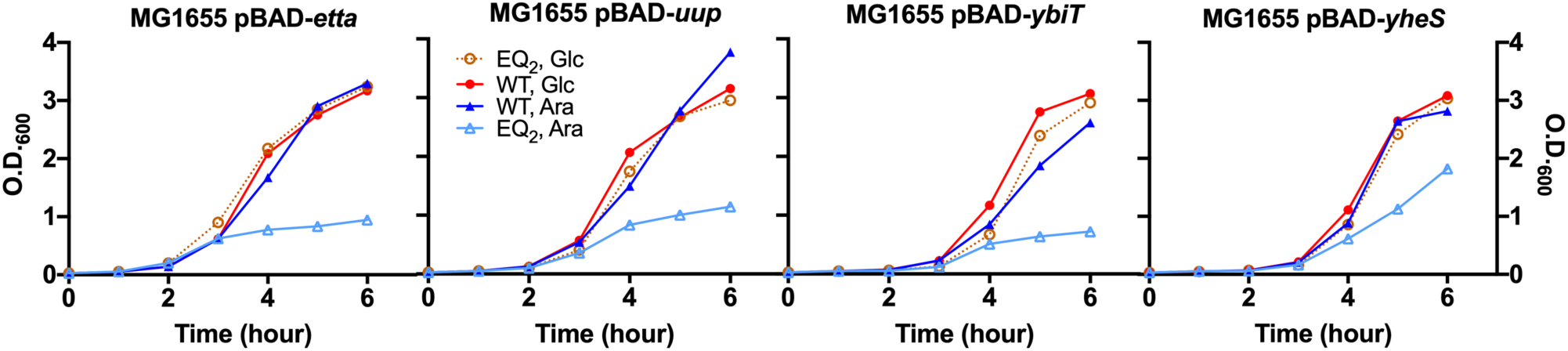
Effects of expressing WT or ATP-hydrolysis-deficient EQ_2_ mutants of the four individual E. coli ABCF paralogs in vivo. Growth at 37 °C was initiated at time zero by diluting overnight cultures 1:100 into fresh unbuffered LB containing 0.4% (w/v) glucose to repress expression or 0.1% (w/v) arabinose or to induce overexpression of the indicated ABCF variants, which were cloned under arabinose promoter control in an ampicillin-resistant pBAD plasmid. The overnight cultures used to initiate the experiment were grown at 37 °C in the same medium with 0.4% (w/v) glucose. This experiment was conducted using *E. coli* strain MG1655^120^.

Similar to previous results ^36^, equivalent *in vivo* expression experiments on EQ_2_ mutants of each of the other three *E. coli* ABCF paralogs demonstrate that they all inhibit cell growth (**Fig. 2**). EQ_2_-YbiT inhibits it as strongly as EQ_2_-EttA, whereas EQ_2_-Uup inhibits it almost as strongly, and EQ_2_-YheS inhibits it significantly but not as strongly as the other paralogs. Expression of WT YbiT produces a much weaker but reproducible inhibition of cell growth, while expression of WT Uup or WT YheS do not (**Fig. 2**), as previously observed for WT EttA ^13^. These results indicate that the ATP-bound conformations of all four *E. coli* ABCF paralogs inhibit some physiological process required for cell growth at 37 °C in LB medium, which is likely to be mRNA translation based on previous observations ^36^ and results presented below and in the accompanying paper ^51^.

### EQ_2_ mutants of the E. coli ABCF paralogs stabilize the GS1 conformation of the ribosome

We previously used single-molecule fluorescence resonance energy transfer (smFRET) experiments in *E. coli* PRE complexes lacking an A-site peptidyl-tRNA (*i.e.*, PRE^−A^ complexes) to demonstrate that, in the presence of ATP, EQ_2_-EttA binds stably to these complexes and stabilizes them in a global conformational state called Global State (GS) 1 or Macro State (MS)-I ^13^, preventing them from spontaneously fluctuating into a different global conformational state called GS2 or MS-II ^52,102,103^. Equivalent smFRET experiments performed on EQ_2_ mutants of all four *E. coli* ABCF paralogs demonstrates that they all interact with ribosomes in a similar manner, although EQ_2_-Uup and EQ_2_-YheS show noteworthy differences in their interaction dynamics (**Fig. 3**).

**Figure 3.**
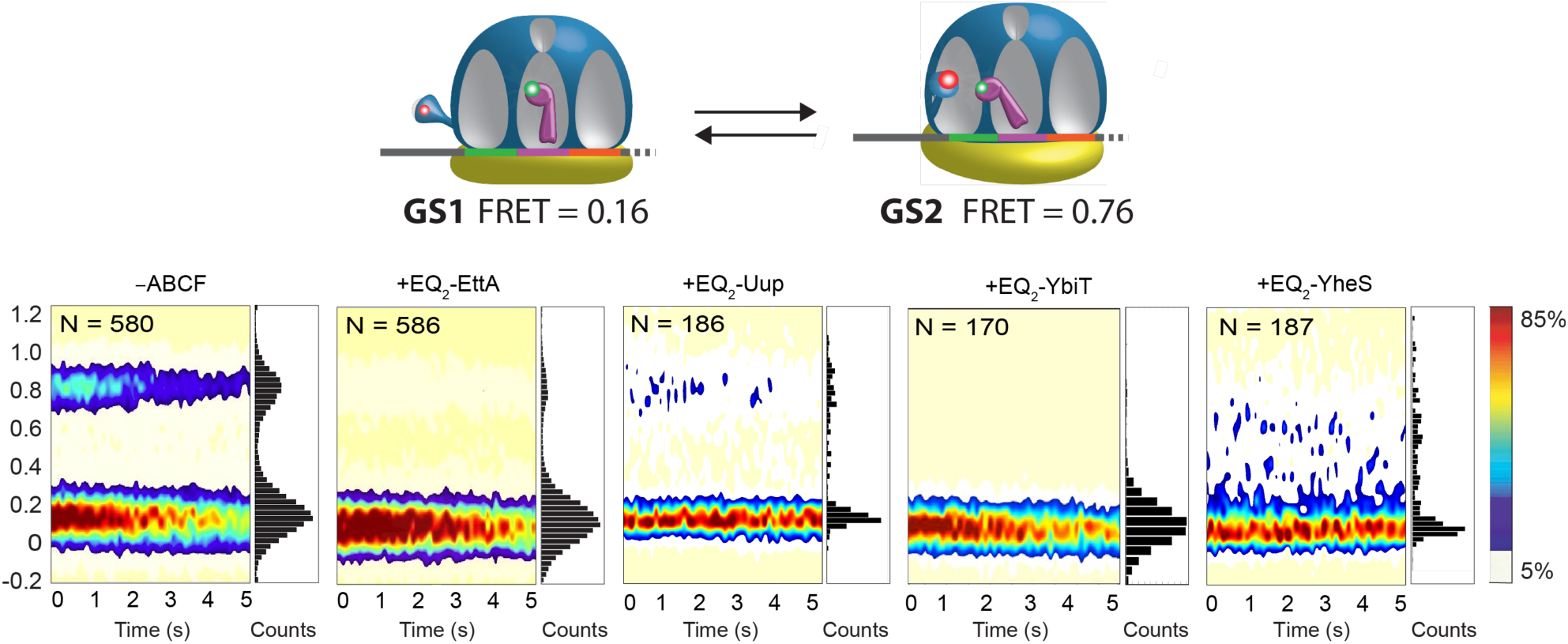
Characterization of the effects the E. coli ABCF paralogs on the conformational dynamics of PRE^−A^ complexes using smFRET. PRE^−A^ complexes bearing a Cy3-labeled, deacylated tRNA^Phe^ in the P site and a Cy5-labeled ribosomal protein uL1 on the apical tip of the uL1 stalk (as schematized at top) were imaged in a buffer containing 2 mM Mg^2+^-ATP using total internal reflection fluorescence (TIRF) microscopy ^103,104,119^. Surface contour plots of the time evolution of FRET are shown for PRE^−A^ complexes in the absence of an ABCF protein (lower left panel) or in the presence of 6 µM EQ_2_-EttA ^12,13^, EQ_2_-Uup, EQ_2_-YbiT, or EQ_2_-YbiT (lower right panels). Contour plots were generated by superimposing individual E_FRET_ *versus* time trajectories. They are colored from white (lowest populated) to red (highest populated), as calibrated by the scale bars on the far right. The bar graphs to the right of each contour plot show a 1D histogram of the E_FRET_ measured during the first second of each experiment. The number of EFRET *versus* time trajectories contributing to each contour plot at time zero, given by the variable N, decreases during the time-course of the measurement due to photo-bleaching of the Cy3 or Cy5 fluorophores. These assays were conducted at 25 °C in Polymix Buffer (100 mM KCl, 5 mM NH_4_OAc, 15 mM Mg(OAc)_2_, 0.5 mM Ca(OAc)_2_, 0.1 mM EDTA, 10 mM 2-mercaptoethanol, 5 mM putrescine dihydrochloride, 1 mM spermidine free base, 50 mM Tris-acetate, pH 7.0).

These smFRET experiments monitor the distance between a Cy3 FRET donor fluorophore attached to a deacylated tRNA^Phe^ bound in the P site of the ribosome and a Cy5 FRET acceptor fluorophore attached to ribosomal protein uL1, a component of the uL1 stalk in the 50S subunit ^104^ (schematic at top of **Fig. 3**). In the absence of any factors bound to the ribosome, this PRE^−A^ complex spontaneously fluctuates between the GS1 and GS2 conformation^52,102,103^. In GS1, the ribosomal subunits are in their ‘non-rotated’ orientation, the Cy3-labeled P-site tRNA is in its ‘classical’ configuration, and the Cy5-labeled uL1 stalk is in its ‘open’ conformation, corresponding to a FRET efficiency (E_FRET_) centered at 0.16^104,105^ (left in schematic at top of **Fig. 3**). In GS2, the ribosomal subunits are in their ‘rotated’ orientation, the Cy3-labeled P-site tRNA is in its ‘hybrid’ configuration, and the Cy5-labeled uL1 stalk is in its ‘closed’ conformation, corresponding an E_FRET_ centered at 0.76 (right in schematic at top of **Fig. 3**). Consequently, spontaneous fluctuations between GS1 and GS2 result in transitions between the 0.16 and 0.76 E_FRET_ values (leftmost “-ABCF” panel at bottom of **Fig. 3**).

This smFRET assay demonstrates that, in the presence of 2 mM ATP, a 6 µM concentration of EQ_2_-Uup greatly reduces the fractional occupancy of GS2, while the same concentration of EQ_2_-EttA, EQ_2_-YbiT, or EQ_2_-YheS blocks detectable occupancy of GS2 (bottom of **Fig. 3**). These results suggest that the EQ_2_ mutants of all four *E. coli* ABCF paralogs bind preferentially to ribosomes in GS1, strongly stabilizing this global conformation and thereby inhibiting transitions into GS2. This inference enables the approximate binding affinities of the EQ_2_-ABCF proteins for the PRE^−A^ complex to be inferred from the reduction in the occupancy of GS2, and this analysis supports the affinities of all four being substantially better than 6 µM. The low level of GS2 occupancy in the presence of EQ_2_-Uup compared to the lack of detectable occupancy in the presence of the other *E. coli* EQ_2_-ABCF proteins supports their binding affinities being significantly stronger than EQ_2_-Uup.

Analyses of cryo-EM structures and population distributions presented in the accompanying paper ^51^ are consistent with these smFRET results. First, the reconstructions produce exclusively conformational classes with the ribosome in GS1 and no classes with it in GS2, consistent with our inference that an EQ_2_-ABCF-bound ribosome is unable to stably sample this global conformational state. Second, assuming that the relative populations of bound *versus* free 70S particle classes reflect the affinities of the EQ_2_-ABCFs for different ribosomal complexes, our cryo-EM data yield affinity estimates for EQ_2_-EttA and EQ_2_-YbiT binding to both a 70S IC and a PRE^−A^ complex and also for EQ_2_-Uup binding to a 70S IC that are all very similar to their affinities for the PRE^−A^ complex inferred from our smFRET experiments.

In contrast, our cryo-EM-based affinity estimates indicate that EQ_2_-YheS has a much weaker ~24 µM affinity for the 70S IC compared to its sub-micromolar affinity for the PRE^−A^ complex inferred from our smFRET experiments (**Fig. 3**). Notably, initiator tRNAs including tRNA^fMet^ contain an additional bulged nucleotide in their elbow region that is not present in most elongator tRNAs ^106–108^, and the stereochemical analyses presented in the accompanying paper ^51^ show direct contacts between all four *E. coli* ABCF paralogs and the elbow region of the P-site tRNA in the 70S IC. In contrast, there are no direct contacts to the formylmethionine residue that is covalently bonded to the 3’-terminus of fMet-tRNA^fMet^. These structural observations suggest that the approximately two order-of-magnitude higher binding affinity of EQ_2_-YheS for a PRE^−A^ complex compared to a 70S IC is more likely to be attributable to the specificity of EQ_2_-YheS for ribosomal complexes harboring an elongator *versus* initiator tRNA^fMet^ in the P site rather than specificity for a deacylated *versus* acylated tRNA in the P site. In any event, the available affinity data indicate that EQ_2_-YheS interacts comparatively weakly with the 70S IC but is likely to interact with high affinity with elongating ribosomal complexes, while EQ_2_-EttA, EQ_2_-YbiT, and EQ_2_-Uup have similarly high affinities for both.

Our smFRET data show PRE^−A^ complexes occasionally fluctuate into GS2 for a period of ~200 milliseconds even when stabilized in the GS1 state by EQ_2_-Uup, but they never reach this state when interacting with EQ_2_-EttA, EQ_2_-YbiT, or EQ_2_-YheS under the same conditions (**Fig. 3**). As discussed above, the simplest explanation for this behavior is that EQ_2_-Uup can dissociate from PRE^−A^ complexes on the second timescale even when trapped in the ATP-bound conformational state by the hydrolysis-blocking EQ_2_ mutations, while the other *E. coli* EQ_2_-ABCF paralogs all dissociate much more slowly when trapped this conformational state. Notably, the ~200 msec lifetime of the GS2 state in the presence of EQ_2_-Uup (**Fig. 3**) is approximately equivalent to its previously established lifetime in the PRE^−A^ complex in the absence of any ribosome-interacting protein ^104^, and this equivalence supports our interpretation that fluctuations into GS2 in the presence of EQ_2_-Uup reflect its dissociation from the PRE^−A^ complex.

In the presence of EQ_2_-YheS, the PRE^−A^ complex shows fluctuations out of GS1 that have a similar ~200 msec lifetime to those observed in the presence of EQ_2_-Uup but that exhibit a distinct range of intermediate E_FRET_ values that are higher than the 0.16 value characteristic of GS1 but lower than the 0.76 value characteristic of GS2 (**Fig. 3**). The failure to reach GS2 in the presence of EQ_2_-YheS indicates this EQ_2_-ABCF is not fully released from the PRE^−A^ during the 5-second experimental observation time. Significant occupancy of states with E_FRET_ values intermediate between GS1 and GS2 are not observed in the presence of the EQ_2_ mutants of any of the other *E. coli* ABCF paralogs. Given the structure of the EQ_2_-YheS-bound 70S IC in GS1 reported in the accompanying paper ^51^, the intermediate E_FRET_ values observed for the EQ_2_-YheS-bound PRE^−A^ complex likely reflect rearrangements of EQ_2_-YheS and the P-site tRNA that bring the uL1 stalk and P-site tRNA closer to each other, and these conformational changes in the E site could potentially be accompanied by changes in global ribosome conformation including intersubunit rotations. While additional research will be required to characterize the structural and functional features of these intermediate states, they demonstrate that EQ_2_-YheS interacts with PRE^−A^ complexes in a significantly different manner than the other *E. coli* ABCF paralogs.

### In vitro tripeptide synthesis reactions show widely varying sensitivity to the EQ_2_ mutants of the four E. coli ABCF proteins

To probe the functional implications of the interactions of the *E. coli* ABCF paralogs with translating ribosomes, we used tripeptide synthesis assays to examine their influence on the first two steps of polypeptide synthesis following initiation of mRNA translation on the *E. coli* 70S IC ^13,109^ (**Fig. 4**). These *in vitro* translation assays evaluate the synthesis of a Met-Phe-Lys tripeptide from a model mRNA. A 70S IC carrying radiolabeled [^35^S]fMet-tRNA^fMet^ in the P site is enzymatically preassembled (left in schematic at top of **Fig. 4**) prior to beginning tripeptide synthesis by the addition of EF-Ts and EF-G along with ternary complexes comprising EF-Tu(GTP)Phe-tRNA^Phe^ and EF-Tu(GTP)Lys-tRNA^Lys^. The ABCF proteins were added at 3 µM concentration in parallel with the EFs and ternary complexes to initiate translation elongation following assembly of the 70S IC (right in schematic at top of **Fig. 4**). Unreacted [^35^S]fMet, [^35^S]fMet-Phe dipeptide, and [^35^S]fMet-Phe-Lys tripeptide were then separated using electrophoretic thin-layer chromatography (eTLC) ^109,110^ and visualized using phosphorimaging (bottom in **Fig. 4**) to evaluate the kinetics of synthesis of the two peptide bonds in the tripeptide.

**Figure 4.**
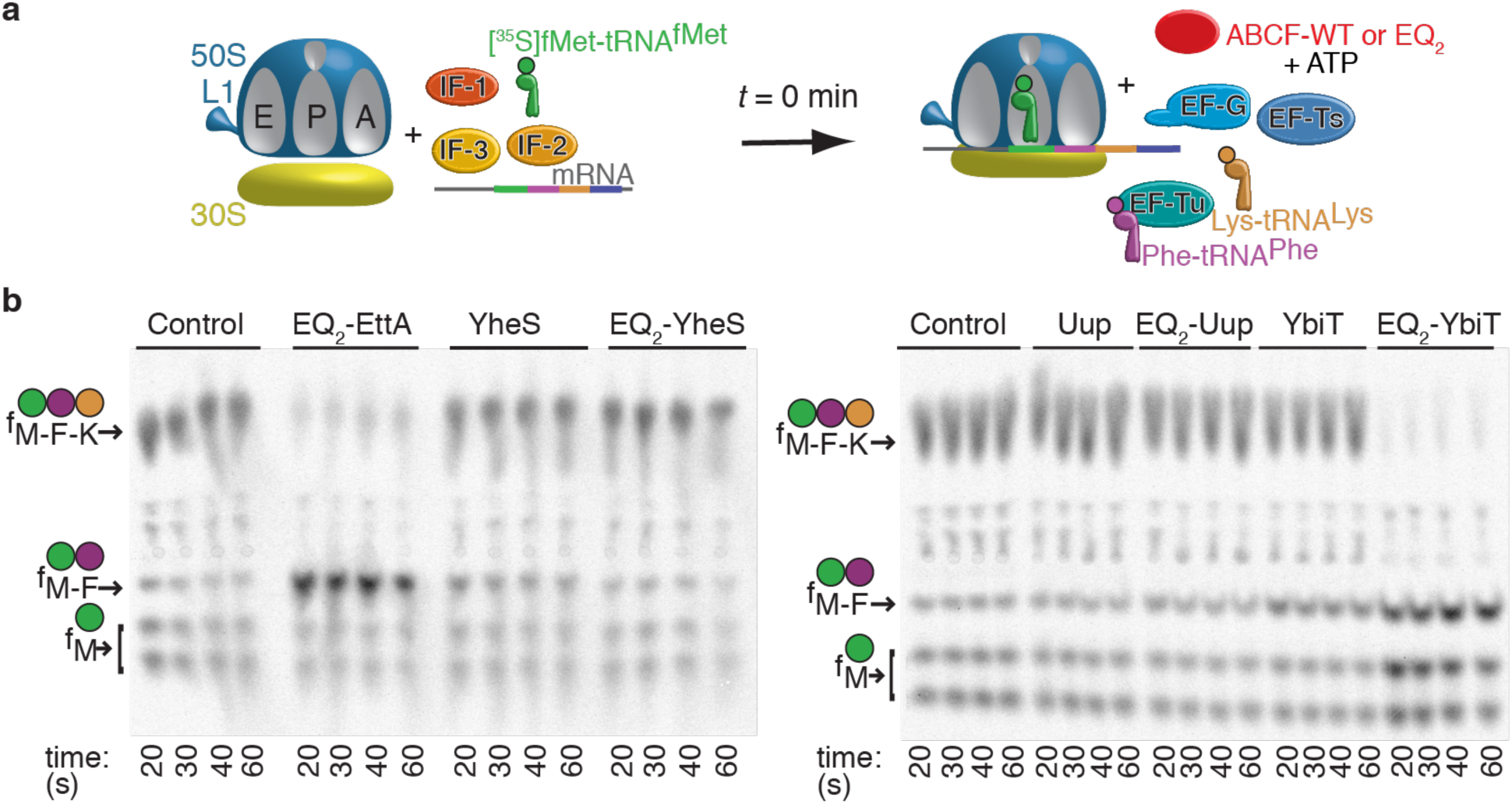
Tripeptide synthesis assays of the influence of purified ABCF paralogs on the initial steps of protein synthesis by E. coli ribosomes in vitro. (a) Schematic diagram of experimental design ^13,110,118^. Polypeptide synthesis cannot proceed beyond the Met-Phe-Lys tripeptide product due to the absence in the *in vitro* translation reactions of an aminolacyl-tRNA cognate to the fourth codon (for Gln) in the pT7gp32.1-20 mRNA template. **(b)** Phosphorimages of eTLC plates after incubation of complete reaction mixtures for the indicated times. The labels on the left indicate the migration positions of f-[^35^S]Met and the f-[^35^S]Met-Phe dipeptide and f-[^35^S]Met-Phe-Lys tripeptide products. The WT ABCF proteins or variants harboring EQ_2_ mutations that trap them in their pre-hydrolysis ATP-bound conformations ^13, 27–32,84,85, 94–101^ were added to a final concentration of 2.9 µM (YheS and Uup variants), 3.4 µM (EttA variants), or 3.5 µM (YbiT variants). Reactions were conducted at 37 °C with 0.3 µM f-[^35^S]Met-tRNA^fMet^ in Polymix Buffer (100 mM KCl, 5 mM NH_4_OAc, 0.5 mM Ca(OAc)_2_, 0.1 mM EDTA, 1 mM spermidine, 5 mM putrescine, 3.5 mM Mg(OAc)_2_, 6 mM 2-mercaptoethanol, 50 mM Tris-OAc, pH 6.9).

These assays demonstrate that EQ_2_-EttA efficiently arrests ribosomes specifically after synthesis of the first peptide bond, producing rapid accumulation of the dipeptide, as previously observed ^13^ (**Fig. 4**). In contrast, EQ_2_-YbiT significantly inhibits synthesis of the first peptide bond and strongly blocks synthesis of the second peptide bond, producing slower accumulation of the dipeptide, while WT-YbiT seems to retard synthesis of the first peptide bond without influencing synthesis of the second peptide bond, producing some transient accumulation of the dipeptide. These observations correlate with the structure of EQ_2_-YbiT bound to the 70S IC reported in the accompanying paper ^51^, where the positioning of the P-site tRNA is not favorable for peptide bond formation. Notably, neither the WT enzymes nor EQ_2_ mutants of Uup or YheS have any clear influence on tripeptide synthesis in these eTLC assays (**Fig. 4**), again highlighting the differences in the functional interactions of the four *E. coli* ABCF proteins with ribosomal complexes. The lack of detectable inhibition of tripeptide synthesis by EQ_2_-YheS is likely attributable to its much lower affinity for the 70S IC ^51^ compared to the PRE^−A^ (**Fig. 3**) and other ribosomal elongation complexes, while the lack of inhibition by EQ_2_-Uup is likely to reflect its relatively facile and rapid exchange from ribosomal complexes, even when locked in the ATP-bound conformation. The Discussion section explains these inferences in detail as well as how the behavior exhibited by the four *E. coli* ABCF paralogs in all of our *in vitro* translation assays is consistent with the affinity data and cryo-EM structures presented here and in the accompanying paper ^51^.

### EQ_2_ mutants of the four E. coli ABCF proteins all inhibit in vitro translation of a reporter mRNA but with widely varying concentration dependence

We also examined the influence of the EQ_2_ mutants of the four *E. coli* ABCF proteins as well as the corresponding WT proteins on the net yield of firefly luciferase from *in vitro* translation reactions conducted using the PURExpress system ^111^, which contains fully defined components (*i.e.*, purified ribosomes, translation factors, synthetases, tRNA, and mRNA). WT EttA, YbiT, and Uup at concentrations up to 9 µM do not significantly alter the yield of luciferase activity, which requires synthesis of the full-length, 550-amino-acid enzyme, while 2 µM WT-Uup produces a small increase in yield that is statistically significant at the 5% confidence level (upper row in **Fig. 5**). In contrast, the EQ_2_ mutants of the four paralogs all strongly inhibit production of luciferase, although the concentrations producing equivalent reductions in yield span a roughly two order-of-magnitude range (lower row in **Fig. 5**). A roughly 50% reduction in luciferase yield is observed in the presence of 0.15 µM EQ_2_-YbiT, 0.5 µM EQ_2_-EttA, 2 µM EQ_2_-YheS, or 9 µM EQ_2_-Uup. Given that these *in vitro* translation reactions were conducted using a 300 nM ribosome concentration, the inhibition produced by EQ_2_-YbiT is nearly stoichiometric, indicating its effective affinity must be far below the ribosome concentration (*i.e.*, in or below the low nanomolar range). The effective affinity for EQ_2_-EttA is higher but must be significantly under 500 nM, while those for YheS and Uup are on the order of 2 µM and 9 µM, respectively. Notably, all 550 residues of luciferase are synthesized with approximately half of the baseline efficiency in the presence of 9 µM Uup even though the smFRET assays indicate it has a binding affinity for the PRE^−A^ complex well below 6 µM (**Fig. 3**). The luciferase synthesis assay results (**Fig. 5**) therefore reinforce the conclusions from the tripeptide synthesis assays (**Fig. 4**), namely that EttA, YbiT, and YheS have relatively straightforward effects on mRNA translation consistent with their observed properties in biophysical assays (**Fig. 3** and the cryo-EM studies in the accompanying paper ^51^), while Uup exhibits more complex behavior addressed in the Discussion section below.

**Figure 5.**
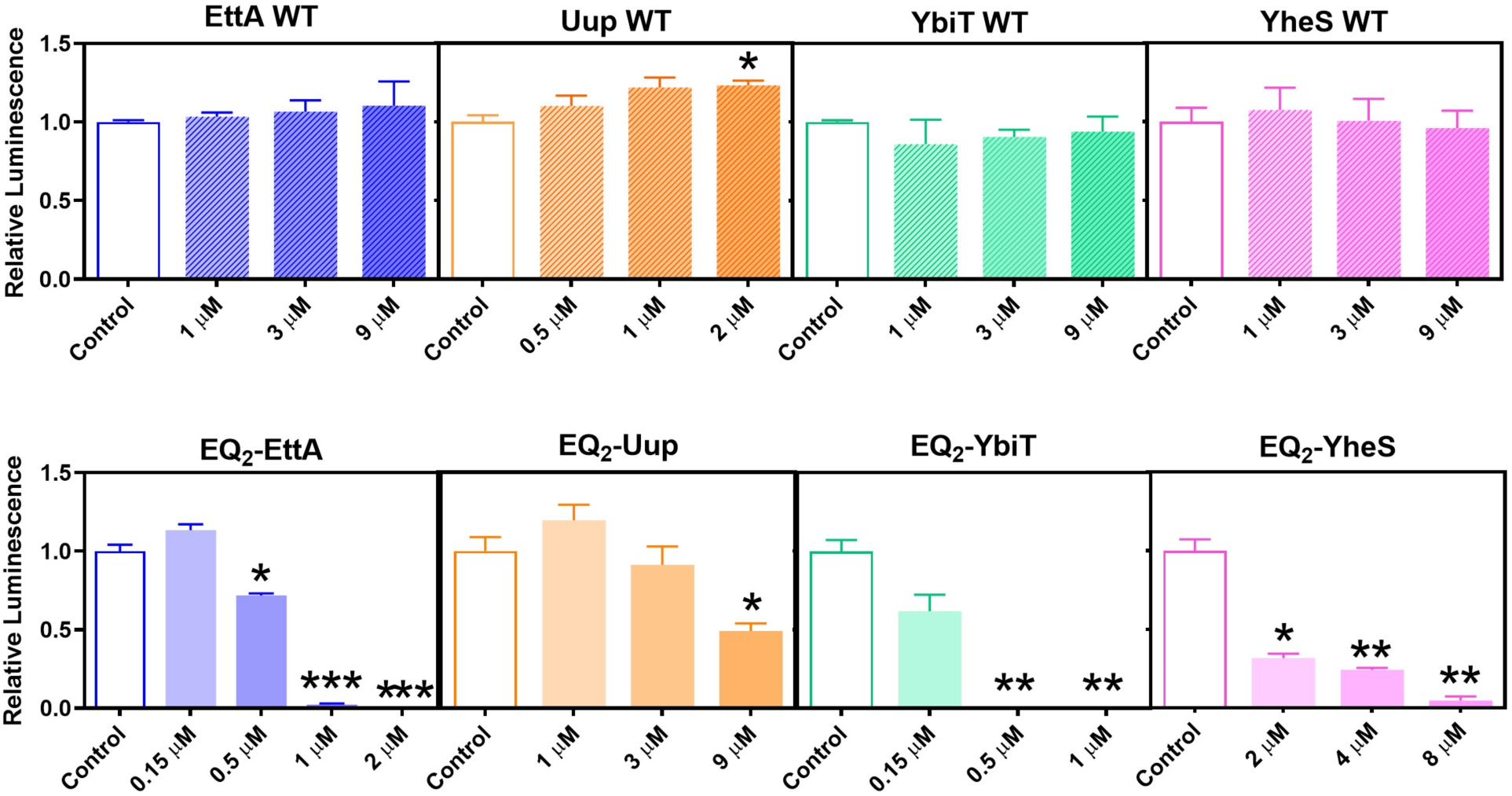
Effects of purified ABCF paralogs on in vitro translation of an mRNA encoding luciferase using a fully defined E. coli translation system. The PURExpress *in vitro* translation system ^111^, which contains purified ribosomes and other translation components, was programmed with purified mRNA encoding firefly luciferase. Purified WT ABCF proteins or variants harboring EQ_2_ mutations that trap them in their pre-hydrolysis ATP-bound conformations ^13, 27–32,84,85, 94–101^ were added at the indicated concentrations. The mRNA was transcribed *in vitro* using bacteriophage T7 RNA polymerase and contains the standard 5’-untranslated region (UTR) produced by the T7 promoter in pET vectors. Luminescence assays were used to measure the yield of active luciferase from *in vitro* translation reactions conducted at 37 °C for 2 hours

## DISCUSSION

The genetic, biochemical, and biophysical data presented in this paper together with the cryo-EM structures reported in the accompanying paper ^51^ demonstrate that the four ABCF paralogs encoded in the genome of *E. coli* K12 (EttA, Uup, YbiT, and YheS) all functionally interact with translating ribosomes, but with distinct biochemical properties (**Figs. 3-5**) and different physiological consequences (**Figs. 1-2**). These results significantly expand understanding of the biochemistry of the ABCF protein family ^7,10,11,112^.

The most mechanistically resolved data on *E. coli* ABCF paralog function is provided by the tripeptide synthesis assays in **Fig. 4**, which dissect the kinetics of synthesis of the first two peptide bonds in a nascent polypeptide after formation of the 70S IC. These experiments were conducted with both the WT enzymes and EQ_2_ mutants that block ATP hydrolysis and trap the enzymes in their ATP-bound, pre-hydrolysis conformations ^84,85, 94–101^. Echoing previous results ^13^, the tripeptide synthesis assays demonstrate that EQ_2_-EttA interacts very strongly with the 70S IC and kinetically traps it after synthesis of the first peptide bond, resulting in stable accumulation of the dipeptide product (**Fig. 4**). These biochemical results are completely consistent both qualitatively and quantitatively with the cryo-EM results presented in the accompanying paper ^51^, which show EQ_2_-EttA bound in the E site of a 70S IC in GS1 and stabilizing the acceptor stem of the P-site tRNA in the proper geometry for peptide bond synthesis in the PTC. Therefore, EQ_2_-EttA should promote synthesis of the first peptide bond in a nascent protein, exactly as observed in the tripeptide synthesis assays. The failure to observe any tripeptide synthesis in these assays is consistent with inhibition of ATP hydrolysis resulting in EQ_2_-EttA being kinetically trapped in the E site of the PRE complex that results following peptide-bond formation, with its binding energy being strong enough to block the subsequent EF-G-mediated tRNA-mRNA translocation. At a similar low-micromolar concentration to that used in the tripeptide synthesis assays, EQ_2_-EttA completely blocks *in vitro* translation of an mRNA encoding full-length luciferase (**Fig. 5**), consistent with its very strong inhibition of tripeptide synthesis (**Fig. 4**). The cryo-EM structures reported in the accompanying paper ^51^ and previous results ^14,53^ demonstrate that EQ_2_-EttA stabilizes either a 70S IC or PRE^−A^ complex in GS1, consistent with its ability to prevent stable transitions to GS2 in the PRE^−A^ smFRET assays reported here (**Fig. 3**).

The behavior of EQ_2_-YbiT is also entirely consistent between the biochemical and biophysical studies reported in this paper and the cryo-EM studies reported in the accompanying paper ^51^. The cryo-EM reconstructions show five conformations of EQ_2_-YbiT, all of which show an equivalent perturbation of the catalytic geometry at the PTC driven by direct interaction of this ABCF protein’s PtIM with the formylmethionylated acceptor stem of the P-site tRNA. The different conformations of EQ_2_-YbiT vary in the orientations of the subdomains in their ABC domains within the E site of a 70S IC or PRE^−A^ complex in GS1. While the different conformations all show two ATP molecules encapsulated between the tandem ABC domains, the ATPase active sites of YbiT adopt a catalytically competent configuration in some of the conformations but not others ^51^. Nonetheless, all of the conformations show an equivalent ~3 Å shift in the location of the terminal 3’-hydroyxyl group in the P-site tRNA and the covalently bonded formylmethionine residue compared to the optimal catalytic geometry observed in reference structures. This stereochemical perturbation is likely to inhibit peptide bond synthesis, and YbiT would presumably have an equivalent effect when a nascent polypeptide chain is covalently bound to the P-site tRNA. Consistent with this inference, EQ_2_-YbiT substantially slows dipeptide synthesis in the tripeptide synthesis assays and completely blocks tripeptide synthesis (**Fig. 4**), with the latter effect likely reflecting ATP-bound EQ_2_-YbiT being kinetically trapped in the E site of the ribosomal complex following peptide-bond formation and blocking subsequent EF-G-mediated tRNA-mRNA translocation. Consistent with these results, a similar low micromolar concentration of EQ_2_-YbiT completely blocks *in vitro* translation of an mRNA encoding full-length luciferase (**Fig. 5**). The stabilization of GS1 that is observed in our smFRET assays with a PRE-A complex (**Fig. 3**) is also consistent with the ribosome conformation observed in our cryo-EM structure of EQ_2_-YbiT bound to a PRE^−A^ complex, which matches the conformation observed when bound to a 70S IC ^51^.

WT-YbiT also produces a small degree of transient dipeptide accumulation in the tripeptide synthesis assays (**Fig. 4**). This effect could be attributable to relatively slow release of YbiT after binding in the E site of the ribosomal complex because occupancy of the non-catalytic conformations of its ATPase active sites observed in the cryo-EM structures retards attainment of the catalytically active conformation. This effect would slow ATP hydrolysis by YbiT, which is required for release from the ribosomal complex, and it could thereby retard EF-G-mediated translocation and continuation of polypeptide synthesis. This effect could also account for the modest inhibition of cell growth produced by overexpression of WT-YbiT in vivo (**Fig. 2**).

The lack of an effect of EQ_2_-YheS in the tripeptide synthesis assays (**Fig. 4**) is also consistent with the cryo-EM studies reported in the accompanying paper, which indicate it has a weak binding affinity for the 70S IC, on the order of ~24 µM ^51^. Given this affinity, the ~3 µM concentration of the protein used in the tripeptide synthesis assays should have a minimal interaction with the 70S IC prior to the initial round of peptide bond synthesis to form the dipeptide and subsequent EF-G-mediated translation. Given the high affinity binding and stable association of EQ_2_-YheS with the PRE^−A^ complex observed in our smFRET studies (**Fig. 3**), the failure to observe any inhibition of tripeptide synthesis suggests that accommodation of the aminoacyl-tRNA bearing the third amino acid in the A site of the ribosomal complex and the subsequent round of peptide bond formation occurs more rapidly than release of the deacylated initiator tRNA^fMet^ from the E site and subsequent binding of EQ_2_-YheS to this vacated site. This inference is consistent with assays showing slow release of deacylated tRNA from the E site at the high Mg^++^ concentration used in the tripeptide synthesis assays ^12,13^.

The inhibition of luciferase synthesis by EQ_2_-YheS (**Fig. 5**) even though it does not inhibit tripeptide synthesis (**Fig. 4**) indicates that it can stop elongating ribosomes at a low micromolar concentration. These results are consistent with the ability of EQ_2_-YheS to stabilize GS1 in our PRE^−A^ smFRET studies (**Fig. 3**), which indicate that the protein has strong affinity for ribosomal complexes carrying a deacylated elongator tRNA in the P site, and the cryo-EM structures of the 70S IC and PRE^−A^ complex presented in the accompanying paper show that EQ_2_-YheS shifts the 3’-hydroyxyl group and covalently bonded formylmethionine on the P-site tRNA away from optimal catalytic geometry in a similar manner to EQ_2_-YbiT, which clearly inhibits translation (**Figs. 4-5**). EQ_2_-YheS would presumably have an equivalent effect perturbing the interaction geometry at the PTC when the P-site tRNA is covalently bonded to a nascent polypeptide chain. These results suggest that EQ_2_-YheS should inhibit elongating ribosomes as long as they have an empty E site in which it can bind.

However, given that the efficiency of luciferase synthesis is assayed based on measurement of enzyme activity, an exceedingly weak inhibitory activity is sufficient to explain the observed result because blocking synthesis of any of its 550 amino acids is sufficient to reduce the yield of full-length, enzymatically active luciferase. In this context, the 75% reduction in yield produced by 2 µM EQ_2_-YheS seems weak compared to the ostensibly complete saturation of the PRE^−A^ complex by a 6 µM concentration of this mutant protein in the smFRET assays (**Fig. 3**), suggesting that most elongation steps during the translation of luciferase are insensitive to EQ_2_-YheS. The relatively weak inhibition of cell growth by EQ_2_-YheS (**Fig. 2**) supports this conclusion. The insensitivity could be attributable to the E site frequently being occupied by a deacylated tRNA during translation elongation, which would sterically block binding of EQ_2_-YheS or any other ABCF protein. Tight coupling of the release of deacylated tRNAs from the E site to EF-G-mediated translocation, which moves the deacylated P-site tRNA into the E site, would lead to continuous occupancy of the E site and blockage of ABCF protein binding in the E site during active translation elongation. More generally, given the cyclical nature of the translation elongation process, even if the E-site tRNA is released prior to EF-G-mediated translocation ^113^, ABCF proteins will only be able to functionally interact with elongating ribosomes if they bind to an empty E site more rapidly than it is refilled by EF-G-mediated translocation of the deacylated tRNA from the P site into the E site.

Therefore, there will be a kinetic competition between ABCF protein binding and translation elongation that is likely to reduce the effective affinity of the ABCF proteins for translating ribosomes compared to any specific ribosomal complex with an empty E site. This effect will tend to attenuate ABCF protein interaction with ribosomal complexes after progression from the initiation stage to the elongation stage of translation, meaning that ABCF proteins will generally interact preferentially with the 70S IC and paused or stalled ribosomal elongation complexes rather than actively elongating ribosomes. The much weaker inhibition of luciferase synthesis by EQ_2_-YheS, which has low affinity for the 70S IC but high affinity for the PRE^−A^ complex, compared to EQ_2_-EttA and EQ_2_-YbiT, which have very high affinity for both, likely reflects a general insensitivity of elongating ribosomes to ABCF protein interaction in the absence a pause or a stall, which derives from the inherent kinetic competition between ABCF protein binding and EF-G-mediated translocation and deacylated tRNA release from the E site.

The behavior of EQ_2_-Uup in our biochemical assays suggests a significant difference in the mechanistic details of its interaction with ribosomal complexes compared to the other *E. coli* ABCF paralogs. EQ_2_-Uup binds to both the 70S IC and the PRE^−A^ complex with ~1 µM affinity based on the smFRET studies reported in this paper (**Fig. 3**) and the cryo-EM studies reported in the accompanying paper ^51^. Its failure to inhibit tripeptide synthesis at an ~3 µM concentration (**Fig. 4**) indicates its interactions with these ribosomal complexes cannot be as kinetically stable as those of the other paralogs. This effect seems unlikely to be attributable to ATP hydrolysis by the protein harboring the EQ_2_ mutations because the cryo-EM structure reported in the accompanying paper ^51^ clearly shows unhydrolyzed ATP in both active sites in the EQ_2_-Uup-bound 70S IC. In any event, ATP-bound EQ_2_-Uup must be released from ribosomal complexes much more rapidly than the EQ_2_ mutants of the other paralogs so that elongation can proceed efficiently despite its binding in the E site of translating ribosomes. Indeed, our smFRET data strongly suggest that EQ_2_-Uup dissociates and reassociates to the E site of PRE^−A^ complexes on a second timescale, while equivalent experiments on the other three paralogs show no evidence of exchange on this timescale. The failure of EQ_2_-Uup to inhibit tripeptide synthesis (**Fig. 4**) is therefore likely to be explained by rapid dissociation from ribosomal complexes on the ~20 second timescale of that experiment, either spontaneously or potentially accelerated by EF-G-mediated translocation. This effect likely contributes to the weak inhibition of luciferase synthesis by EQ_2_-Uup (**Fig. 5**), although its ability to significantly attenuate luciferase synthesis suggests that it released more slowly from a specific conformation, or conformations, of the elongating ribosome that arise during translation of this 550-residue protein that are not significantly occupied during translation of the model tripeptide.

The great phylogenetic prevalence and diversity of the ABCF protein family suggest that its members perform important biological functions. However, the details of these functions remain minimally defined even though the three human paralogs have been implicated in disease-related processes ranging from modulation of immune responses ^37–40,44,114^ to the development and treatment of cancers ^41–49^. Given their involvement in these critical biological processes, it is important to understand the nature and range of the biochemical activities performed by ABCF paralogs. The experimental results on the four *E. coli* ABCF paralogs reported in this paper and the accompanying paper ^51^ reinforce and extend recent research suggesting that ABCF proteins perform diverse biochemical functions related to mRNA translation ^8,12,13, 25–28,59,61,62^. All paralogs characterized to date bind to ribosomal complexes with an empty E site and control the interaction geometry of the aminoacylated/peptidylated acceptor stem of the P-site tRNA with the PTC ^12,14,30,51^. This common mode-of-action suggests different ABCF paralogs may tend to have some degree of overlap in their physiological activities, as we demonstrate for EttA and YbiT in our genetic assays (**Fig. 1c**). However, the results reported in this paper demonstrate that the activities of different paralogs are tuned in distinct ways that are likely to reflect substantial differences in biological function. At least one of the four *E. coli* ABCF paralogs behaves qualitatively differently in every one of the genetic (**Fig. 1**), physiological (**Fig. 2**), biophysical (**Fig. 3**) and/or biochemical (**Figs. 4-5**) experiments reported in this paper. These observations indicate the paralogs have significantly different biochemical and physiological functions despite their homologous structures and common mode of binding to ribosomal complexes ^51^. The function of EttA is understood in considerable detail (ref. ^13^ and manuscripts in preparation^115,116^, and the activity of YbiT seems to be influenced by the presence *vs.* absence of the sub-stoichiometric protein bL33 in the 50S subunit ^51^. However, additional research will be needed to elucidate the biological function of YbiT as well as the functions of Uup, YheS, and the extensive number of completely uncharacterized ABCF paralog groups ^36^. Our studies of the four *E. coli* ABCF paralogs suggest that there is still a substantial amount of functionally uncharacterized biological “dark matter” involved in controlling mRNA translation.

## Author Contributions

G.B., R.L.G. and J.F.H. designed the research and wrote the paper in consultation with the other authors. G.B. and K.-H.W. performed the *in vivo* toxicity assays. G.B., S.S, L.Z., and M.J.N. purified the proteins. S.S., G.B., and K.-H.W. performed the luciferase translation assays. J.F. and N.A.B. performed the smFRET experiments. G.B. and L.Z. performed the tripeptide synthesis assays. G.B., C.S., and F.O. performed the fitness experiments.

## Abbreviations

ABC: ATP-binding cassette
ARE: antibiotic resistance
A site: aminoacyl-tRNA binding site on the ribosome
cryo-EM: cryogenic electron microscopy
EQ_2_: mutations converting both catalytic glutamates in an ABCF protein to glutamine
E site: tRNA exit site on the ribosome
IC: initiation complex
OAc: acetate ion (CH_3_COO^−^)
P site: peptidyl-tRNA binding site on the ribosome
PRE: ribosome in the pretranslocation conformation
PRE^−A^: ribosome in the pretranslocation conformation without a tRNA bound in the A site
PTC: peptidyl-transferase center
PtIM: P-site tRNA Interaction Motif
TCEP: tris(carboxyethyl)phosphine
WT: wild type

## Acknowledgements

We thank J. Boothe for assistance with protein purification, L. Dietrich for use of equipment, E. Bailey for providing constructs for the luciferase assays, and the members of the Hunt and Gonzalez laboratories for advice. This work was supported by grants from the National Institute of General Medical Sciences of the U.S. National Institutes of Health to the Northeast Structural Genomics Consortium (GM074958), J.F.H. (GM127883), and R.L.G. (GM084288 and GM137608) and by a grant from the U.S. National Science Foundation to R.L.G. (MCB0644262). G.B. and F.O. were supported by funds from the CNRS (UMR8261), Paris Cité University, the LABEX program (DYNAMO ANR-11-LABX-0011), and two ANR grants (EZOtrad/ANR-14-ACHN-0027 and ABC-F_AB/ANR-18-CE35-0010). F.O. received a fellowship from the Edmond de Rothschild Foundation.

## METHODS

### Competitive fitness assays

Assays were conducted in unbuffered LB (USB-Affymetrix) at 37 °C using the methods we previously used to characterize the Δ*ettA* strain ^13^. Clean mutant strains harboring a complete knockout of each individual ABCF gene or all four simultaneously were constructed from the sequenced WT MG1655 strain using the λ_red_ recombination method ^82^. Mutagenesis employed PCR products amplified from genomic DNA from Keio Collection strains ^81^ in which the entirety of the coding sequence of the individual *uup* (JW0932-1), *ybiT* (JW0804-1), or *yheS* (JW3315-1) gene is replaced with a kanamycin resistance gene. The amplified DNA segments, which included 400 base pairs upstream and downstream of the deleted ABCF gene, were introduced via electroporation into MG1655 cells harboring a plasmid expressing the λ_red_ recombinase and Gam proteins. Following selection of kanamycin-resistant ABCF knockout strains, the kanamycin resistance gene was removed from the chromosome as previously described ^82^. This gene knockout method is designed to limit polar effects ^81^, but it could alter the expression levels of the products of co-cistronically genes encoded downstream (*yheT*, *yheU*, and *prkB* for *yheS*) or even possibly upstream (*rlmL* for *uup*) of the deleted target gene. Complemented strains of the individual knockouts were constructed using the clonetegration recombination method ^117^ to insert the individual ABCF genes with their native promotors at the P21 genomic locus.

### In vivo expression of WT proteins and EQ_2_ mutants

The wild-type *E. coli* ABCF proteins and variants harboring EQ_2_ mutations in their catalytic bases were cloned with or without a hexahistidine affinity tag into a pBAD vector under arabinose-promoter control. For YbiT, a C-terminal tag with sequence LEHHHHHH was fused in-frame after the final residue in the native protein. For the other proteins, an N-terminal tag was fused in-frame in front of the native initiation codon. A tag with sequence MAHHHHHH was used for EttA and YheS, with EttA’s native initiator GUG codon converted to AUG. A TEV-protease-cleavable tag with sequence MAHHHHHHENLYFQ was used for Uup.

The EQ_2_ mutations were introduced sequentially into each protein using QuikChange II Site-Directed Mutagenesis (Agilent Technologies) with all growth media supplemented with 0.4% (w/v) glucose to mediate catabolite repression of ABCF protein expression from the arabinose promoter. Plasmids were transformed into *E. coli* strain MG1655 for cell growth experiments in unbuffered LB medium in the presence of 100 µg/ml ampicillin at 37 °C in 24 mm glass tubes with moderate aeration. At time zero, overnight cultures in the same medium containing 0.4% (w/v) glucose to maintain catabolite repression of the arabinose promoter were diluted 1:100 into 5 ml of fresh medium containing either the same concentration of glucose or 0.1% (w/v) arabinose to induce protein expression.

### Protein purification

The same plasmids and host strain used for the *in vivo* expression experiments were used for preparative scale protein production. Overnight starter cultures in LB with 0.4% (w/v) glucose were diluted 1:100 into 2-3 L of fresh medium for WT proteins. BL21-LobSTR strain was used for the expression of the EQ_2_ mutants. Overnight starter cultures in LB with 0.4% (w/v) glucose were diluted 1:100 into 3-4 L of fresh Terrific Broth/Amp for the EQ_2_ mutant proteins. These cultures were grown at 37 °C until OD_600nm_ reached ~0.6 for WT or ~2.0 for EQ_2_ mutants, prior to induction of protein expression for 2-3 hours at the same temperature using 0.1% (w/v) arabinose. The proteins were purified as previously described for EttA using Ni-NTA affinity chromatography followed by gel-filtration chromatography in 150 mM NaCl, 5% (v/v) glycerol, 1 mM TCEP, 20 mM Tris-HCl, pH 7.5. Proteins were snap-frozen in small aliquots after concentrating them in this buffer using a Centricon 50 (EMD Millipore, Billerica, MA). Protein concentrations were determined using BSA standards on a Coomassie-Blue-stained SDS-PAGE gel.

### Luciferase in vitro translation assays

Methods were equivalent to those reported earlier. In brief, reactions were conducted at 37 °C using the PURExpress System (New England Biolabs, Ipswich, MA) ^111^ in the absence or presence of increasing concentrations of ABCF protein. Reactions were run for 2 hours using 6 μl of Solution A, 1.8 μl of Factor Mix, 1 μl of T7-luc mRNA at 3 μg/μl, and 5.9 μl of ABCF protein stock or control buffer in a total reaction volume of 15 μl. Ribosome concentration was kept at ~300 nM in the reactions. Luminescence assays were conducted on a Synergy™ 4 microplate reader (BioTek, Winooski, VT) using 3 µl of the *in vitro* translation reaction mixture in 150 µl of luciferase assay reagent (Promega, Fitchburg, WI). The graphs in **Fig. 5** show data from a set of triplicate assays on the WT and EQ_2_ variants.

### Tripeptide synthesis assays

Methods were identical to those reported earlier by Boël *et al.* ^13^ using a minimum *in vitro* translation system ^110,118^ with the 70S IC formed prior to the addition of the ABCF protein in parallel with EFs and ternary complexes. The unreacted f-[^35^S]Met and peptide products were separated using eTLC ^110,118^.

### smFRET assays

As previously described ^103,104,119^, a total internal reflection fluorescence (TIRF) microscope was used to characterize the conformational dynamics of PRE^−A^ complexes assembled on a 5’-biotinylated model mRNA that was used to tether the complexes to the polyethylene glycol-passivated, streptavidin-derivatized surface of a quartz microfluidic flow cell. The buffer described in the figure legend also contained an oxygen-scavenging system comprising 25 nM protocatechuate 3,4-dioxygenase, 2.5mM 3,4-dihydroxybenzoic acid, and 1% (w/v) β-D-glucose. Data were acquired using a 50 millisecond integration time.

